# Cadmium exposure inhibits branching morphogenesis and causes alterations consistent with HIF-1α inhibition in human primary breast organoids

**DOI:** 10.1101/208157

**Authors:** Sabrina A. Rocco, Lada Koneva, Lauren Y. M. Middleton, Tasha Thong, Sumeet Solanki, Sarah Karram, Kowit Nambunmee, Craig Harris, Laura S. Rozek, Maureen A. Sartor, Yatrik M. Shah, Justin A. Colacino

## Abstract

**Background:** Developmental cadmium exposure *in vivo* disrupts mammary gland differentiation, while exposure of breast cell lines to cadmium causes invasion consistent with the epithelial-mesenchymal transition (EMT). The effects of cadmium on normal human breast stem cell development have not been measured.

**Objective:** The objective of this study was to quantify the effects of cadmium exposure on normal breast stem cell proliferation and differentiation.

**Methods:** We tested the effects of two physiologically relevant doses of cadmium: 0.25µM and 2.5μM on reduction mammoplasty patient-derived breast cells using the mammosphere assay, organoid formation in 3D hydrogels, and tested for molecular alterations using RNA-seq. We functionally validated our RNA-seq findings with a HIF-1α transcription factor activity reporter line and pharmaceutical inhibition of HIF-1α in mammosphere and organoid formation assays.

**Results:** 2.5μM cadmium reduced primary and secondary mammosphere formation and branching structure organoid formation rates by 33%, 40%, and 83%, respectively. Despite no changes in mammosphere formation, 0.25μM cadmium treatment inhibited branching organoid formation in hydrogels by 68%. RNA-seq revealed that cadmium treatment downregulated genes associated with extracellular matrix formation and EMT, while upregulating genes associated with metal response including metallothioneins and zinc transporters. In the RNA-seq data, cadmium treatment also downregulated HIF-1α target genes including *LOXL2*, *ZEB1*, and *VIM*. Cadmium treatment significantly inhibited HIF-1α activity in a luciferase assay, and the HIF-1α inhibitor acriflavine ablated mammosphere and organoid formation.

**Discussion:** These findings show that cadmium, at doses relevant to human exposure, inhibited human mammary gland development, potentially through disruption of HIF-1α activity. These findings do not support cadmium being a breast cancer initiator via induction of stem cell proliferation, but instead implicate cadmium as an inhibitor of mammary gland morphogenesis.

## INTRODUCTION

Breast cancer is the most common cancer in women, with an estimated 246,660 incident cases in 2016 in the United States alone (Siegel et al. 2016). Approximately 90% of breast cancers are thought to be due to sporadic, rather than hereditary, alterations, potentially arising due to environmental exposures (Rizzolo et al. 2011). Despite decades of research, the environmental risk factors for breast cancer are still not well understood. The toxic heavy metal cadmium is a naturally occurring known human carcinogen which has been strongly linked to lung cancer through occupational health studies (Waalkes 2003). Its role in breast cancer remains controversial. Multiple case-control studies have reported that urinary cadmium concentrations, a biomarker of cadmium exposure, are higher in breast cancer cases relative to controls (Gallagher et al. 2010; McElroy et al. 2006; Nagata et al. 2013; Strumylaite et al. 2014). Results deriving from prospective cohort studies, are equivocal with some finding positive relationships between cadmium exposure and breast cancer (Julin et al. 2012) and others identifying null associations (Adams et al. 2012; Adams et al. 2014; Eriksen et al. 2014; Eriksen et al. 2017; Sawada et al. 2012).

Experimental work *in vivo* and *in vitro* show that cadmium exposure can alter normal mammary gland development and related developmental pathways. The breast cancer windows of susceptibility hypothesis states that environmental exposures during key developmental time points, particularly *in utero*, during puberty or pregnancy, can disproportionately increase breast cancer risk later in life (Russo 2016). These windows represent times when the mammary gland is undergoing substantial remodeling, driven by a population of proliferating and differentiating stem cells (Visvader and Stingl 2014). Environmental exposures during these time points could alter normal breast stem cell self-renewal, modifying the number of stem cells in the tissue (Ginestier and Wicha 2007), or otherwise influence differentiation pathways, leading to breast cancer or other forms of breast toxicity. The *in vivo* data of cadmium exposure during windows of susceptibility show that life stage of exposure is important, where *in utero* exposures to cadmium display an estrogenic effect leading to an increase in terminal end bud structures (Alonso-Gonzalez et al. 2007; Johnson et al. 2003), while exposures in puberty or adulthood lead to stunted mammary gland development (Davis et al. 2013; Ohrvik et al. 2006). A more limited number of *in vitro* studies of the effects of cadmium in immortalized breast cell lines highlight that cadmium exposure can lead to the acquisition of an invasive phenotype (Benbrahim-Tallaa et al. 2009; Wei and Shaikh 2017), consistent with dysregulation of the important developmental pathway the epithelial-mesenchymal transition (EMT). Despite a growing body of experimental and epidemiological evidence linking cadmium to altered mammary gland development and potentially breast cancer, very little is known about the effects of cadmium exposure on primary non-transformed and non-immortalized human breast stem cells. This is an important research gap as breast stem cells are likely a key target for cadmium’s effects on breast cancer or altered mammary gland development.

The goal of this study was to test the hypothesis that cadmium induces changes in primary human breast stem cells consistent with alterations in stem cell self-renewal and developmental pathways such at EMT. We established a novel model of the effects of environmental exposures during human breast development, integrating an established technique, the mammosphere formation assay (Dontu et al. 2003) with novel 3D breast organoid culture (Sokol et al. 2016). Using functional and high-throughput molecular assays, we identify that cadmium, at doses relevant to human exposure, induces significant alterations in human breast stem cell differentiation and development.

## METHODS

### Human Tissue Procurement

Non-pathogenic breast tissue was isolated from women undergoing voluntary reduction mammoplasty at the University of Michigan. Breast tissue was mechanically and enzymatically digested as previously described (Colacino et al. 2016; Dontu et al. 2003) to yield a viable single cell suspension of human mammary cells. This study was reviewed and approved by the University of Michigan Institutional Review Board (HUM00042409).

### Cadmium Dosing and Mammosphere Formation

A general outline of the experimental design is presented in Figure 1. In previous studies, chronic exposure to 2.5μM cadmium led to transformation of MCF10A cells into cells showing a mesenchymal phenotype with increased invasive capabilities consistent with cancer development (Benbrahim-Tallaa et al. 2009). Studies of cadmium in breast tissue, from breast tumors or benign breast tissues, found mean tissue concentrations of cadmium ranging from 17.5 ng/g to 37 ng/g (Strumylaite et al. 2008; Strumylaite et al. 2011). Assuming a tissue density equal to that of water, these concentrations would correspond to a dose of 0.156 and 0.33μM, respectively. Based on these previously published results and our dose-response experiments, we chose two relevant doses of cadmium for further study, 0.25μM and 2.5μM. The primary mammosphere formation assay is a surrogate readout of breast stem cell activity, while the secondary mammosphere formation assay is a readout of breast stem cell self-renewal capacity (Dontu et al. 2003). Single primary human breast cells were plated in mammosphere formation conditions, in the presence of 0.25μM or 2.5μM cadmium chloride or vehicle control following our previously established protocol (Colacino et al. 2016). Briefly, cells were plated at a concentration of 100,000/mL in ultralow attachment plates (Corning) conditions in MammoCult (StemCell) media. Additional media (either control or cadmium containing) was added at 3 days. Primary mammospheres formed for 7 days, at which point mammospheres larger than 40μm were counted manually. For quantification of secondary mammosphere formation, primary mammospheres were dissociated back into a single cell suspension with TrypLE Express (Life Technologies), counted and resuspended in MammoCult, and plated in non-adherent plates at 100,000 cells/mL. The number of secondary mammospheres formed were quantified manually after 7 days. At least three technical replicates were quantified per condition across six independent biological replicates.

**Figure 1.**
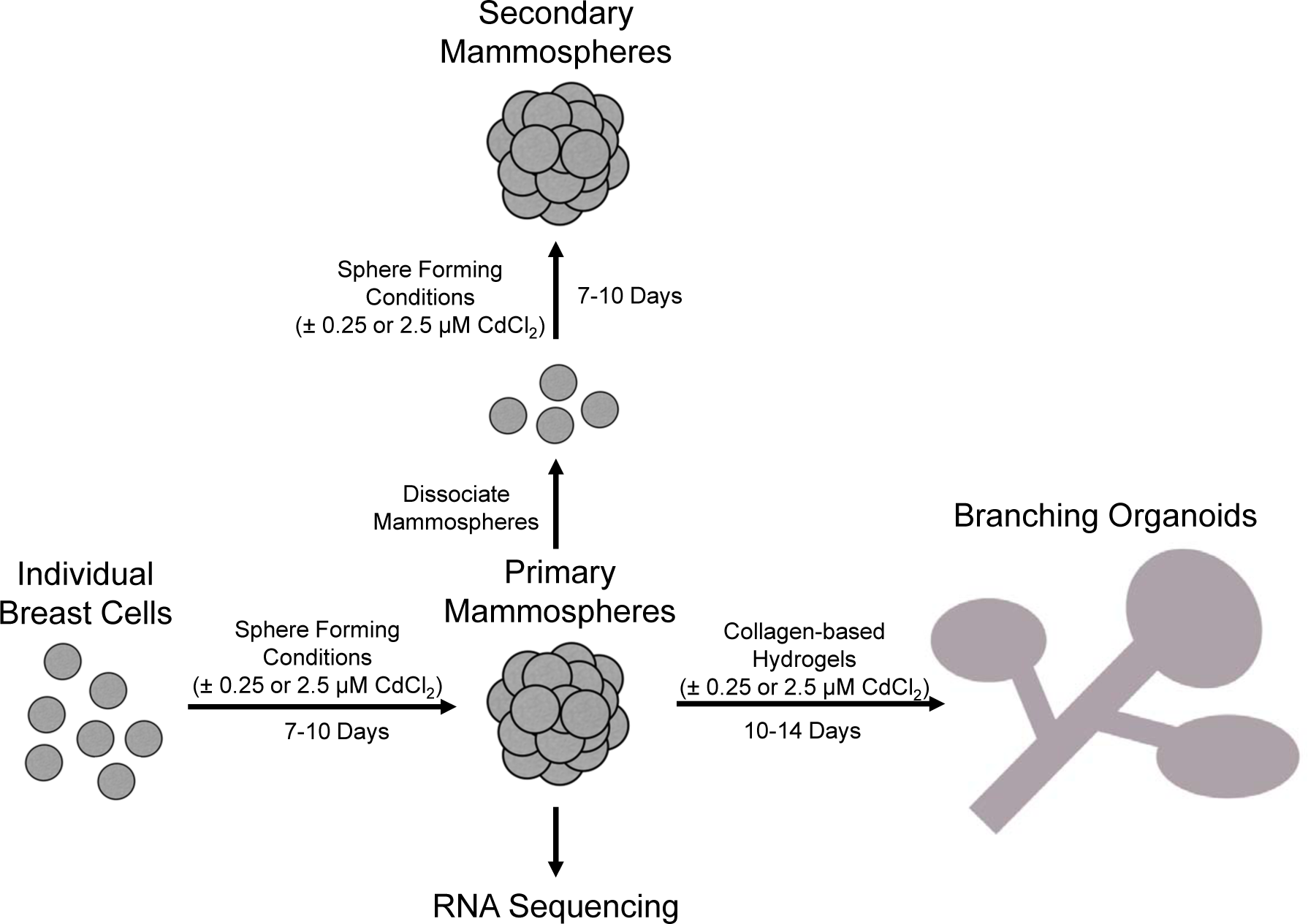
Conceptual diagram of the experimental design. Epithelial cells isolated from voluntary reduction mammoplasty tissues were cultured in the presence or absence of two doses of cadmium in mammosphere forming conditions. Primary mammospheres were either dissociated into single cells and replated in secondary mammosphere formation conditions, plated in collagen-based hydrogels to induce breast organoid formation, or RNA was extracted and sequenced.

### Mammary Organoid Formation

At the end of the 7-10 days of primary mammosphere formation in the presence or absence of cadmium, mammospheres were embedded in floating 3D collagen-based hydrogel scaffolds to induce branching morphogenesis, a process which models mammary gland development. Hydrogels were prepared as previously described (Sokol et al. 2016). Briefly, Laminin (Roche), Fibronectin (Roche), 100 KDa and 700 KDa Sodium Hyaluronate (Lifecore Biomedical), MEGM growth media (Lonza), and primary mammospheres were added to rat tail Collagen Type I (EMD Millipore). We added 0.2M sodium hydroxide (Sigma-Aldrich) to induce hydrogel polymerization, and plated the gels in 24-well plates for 1 hour at 5%CO_2_ and 37°C. After polymerization, hydrogels were detached from the plate using a metal spatula and media was added. The hydrogels were plated in either MEGM media as a control, or MEGM containing 0.25μM or 2.5μM cadmium chloride. These were cultured at 37°C at 5% CO_2_ for 2-3 weeks with media changes every 3-4 days. The number of branching structures that formed in each gel was quantified manually after 10 to 14 days. At least three technical replicates were quantified per condition across at least three independent biological replicates.

### RNA Sequencing of Primary Mammospheres

After 7 days of growth in primary mammosphere formation conditions, we extracted RNA from the cadmium or control treated mammospheres using the AllPrep DNA/RNA micro kit (Qiagen), including an on-column DNase treatment for RNA extraction. RNA concentration and quality was determined using a Nanodrop (Thermo) and Bioanalyzer (Agilent). We depleted ribosomal RNAs with Ribominus (Life) and prepared sequencing libraries utilizing the SMARTer Stranded RNA-Seq kit (Clontech) following the manufacturer’s recommended protocol. Libraries were multiplexed (6 per lane) and sequenced using paired end 50 cycle reads on a HiSeq 4000 (Illumina). Library preparation and sequencing took place at the University of Michigan DNA Sequencing Core Facility following their standard protocols.

### RNA-Seq Data Analysis

The RNA-seq libraries were aligned to the GRCh38.p10 human genome (GRCh38_GencodeV26 <u>https://www.gencodegenes.org/releases/current.html).</u> Quality control of raw fastq files was performed using FastQC v 0.11.5 (Andrews 2012) and MultiQC v0.9 (Ewels et al. 2016) to identify features of the data that may indicate quality problems (e.g., low quality scores, over-represented sequences, and inappropriate GC content). Mapping of raw sequences were performed using STAR-2.5.3.a (Dobin et al. 2013) with default parameters. The RNA-seq library sizes ranged from 25.4 to 66.7 million reads (average 46.3 million) an alignment rate is 66.5%-75.1% (average 71.2%). MultiQC, RSeQC v2.6.4 (Wang et al. 2012) and QoRTS v1.2.26 (Hartley and Mullikin 2015) were used for a second round of quality control (post-alignment), to ensure that only high quality data would be input to expression quantitation and differential expression analysis. Gene expression levels were quantified using Subread v 1.5.2 package FeatureCounts, performing strand-specific pair-end reads counting (Liao et al. 2014).

Normalization and transforming read counts for diagnostic plots, including PCA and hierarchical clustering, were performed using DESeq2 v1.14.1 Bioconductor package (Love et al. 2014). The differential expression testing was conducted with the Bioconductor package edgeR v 3.16.5 using edgeR-robust and the quasi-likelihood functionality of edgeR (Chen et al. 2016; Robinson et al. 2010; Zhou et al. 2014). We conducted filtering of the low expressed genes keeping only genes that are expressed at least 1 CPM (counts per million reads) in at least 3 samples. The comparisons were performed for the following groups: 1) cells treated with 2.5μM cadmium chloride versus control and 2) cells treated with 0.25μM cadmium chloride versus control. P-value adjustments for multiple testing were made using the Benjamini-Hochberg False Discovery Rate approach. Genes and transcripts satisfying criteria (FDR<0.10 and absolute value of fold change |FC| >1.5) were considered differentially expressed (DEGs).

For the set of genes from the comparison 2.5μM cadmium chloride versus control (141 genes with FDR < 0.1) the heatmap visualization with hierarchical clustering with complete linkage and dendrogams by genes and samples were performed to identify the main expression profiles observed across the two treatment conditions and control. Normalized by DESeq2 read counts were log-transformed with pseudo-count (=1) and then data were normalized again by subtracting the overall average expression of each gene from each expression value.

### Pathway Analysis

Enriched Gene Ontology (GO) terms, KEGG pathways and transcription factors for each of the analyzed comparisons were tested using RNA-Enrich (Lee et al. 2016) (http://lrpath-db.med.umich.edu/). RNA-Enrich tests for gene sets that have higher significance values (e.g., for differential expression) than expected at random. By not requiring a cutoff for significance, RNA-Enrich is able to detect both pathways with a few very significant genes and pathways with many only moderate differentially expressed genes. A directional RNA-Enrich test, which tests for significantly up- versus down- regulated gene sets, was run for each comparison using default settings. Only concepts with less than 500 genes were considered for this analysis. Custom code was implemented to reduce redundancy (remove less significant, closely related GO terms) for presenting the top enriched terms by cadmium treatment.

### HIF-1α Inhibition Experiments

Embryonic kidney cell line HEK293T were seeded into a 24-well plate at a cell density of 5 × 104 cells per well. HIF1α response element driven luciferase was co-transfected with β-galactosidase into cells with polyethylenimine (PEI; Polysciences Inc., Warrington, PA) for 24 hours. After 24 hours of transfection, cells were treated with concentrations of 0.25μM, 2.5μM and 25μM cadmium chloride in the presence or absence of FG4592 (100μM, hypoxia-inducible factor (HIF) prolyl hydroxylase inhibitor) for another 24 hours. Cells were lysed in reporter lysis buffer (Promega, Madison, WI) and luciferase assay was performed and normalized to β-galactosidase activity. Primary human breast epithelial cells grown in mammosphere forming conditions were also treated with the pharmaceutical inhibitor of HIF-1α activity acriflavine at concentrations of 1μM and 5μM. Primary and secondary mammosphere formation rates and organoid formation were measured as described above.

### Statistical Analysis

Differences of primary mammospheres, secondary mammospheres, and hydrogel structures were analyzed within individuals using a paired t-test for control to 0.25μM cadmium chloride, control to 2.5μM cadmium chloride, and 0.25μM cadmium chloride to 2.5μM cadmium chloride, or control to either concentration of acriflavine. Totals across all biological replicates were also analyzed for values averaged across all technical replicates using a paired t-test. For all tests, a p-value of 0.05 was considered statistically significant.

## RESULTS

### Effects of Cadmium Exposure on Primary and Secondary Mammosphere Formation

To quantify the effects of cadmium on normal breast stem cell activity and self-renewal capacity, human mammary cells were plated in mammosphere forming conditions in the presence or absence of the two cadmium doses. Across six independent biological replicates, we did not observe a significant alteration in primary mammosphere formation at the lower cadmium dose, but 2.5μM cadmium chloride exposure decreased primary sphere formation by approximately 33% (Figure 2A). Similar results were observed for secondary sphere formation, where 0.25μM cadmium chloride had no significant effect and the 2.5μM dose inhibited secondary mammosphere formation by approximately 40% (Figure 2C). Representative images of the mammospheres formed in the various experimental conditions are displayed in Figure 2B and 2D.

**Figure 2.**
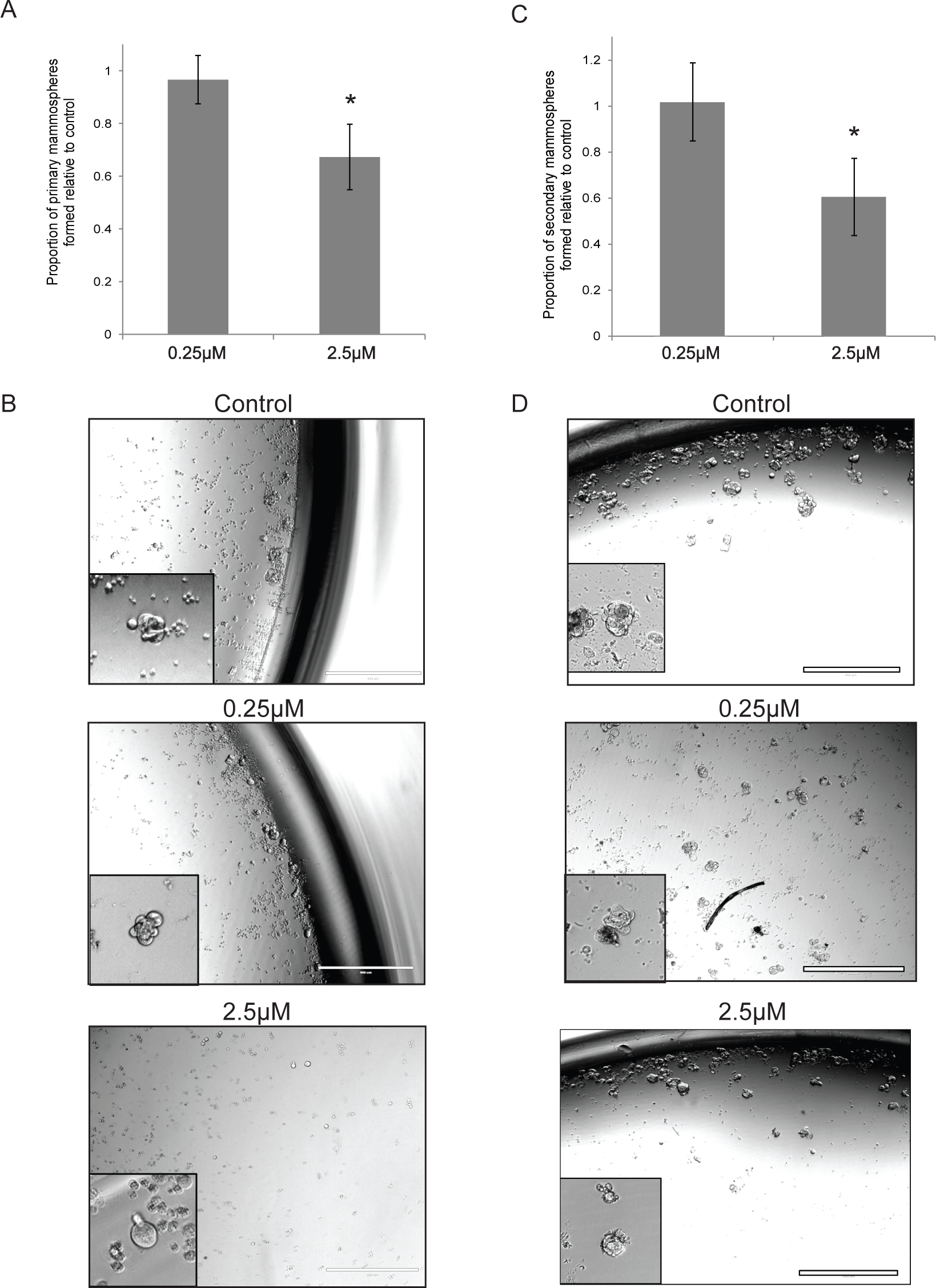
The effects of cadmium on mammosphere formation. (A) The effects of 0.25μM or 2.5μM cadmium chloride on primary mammosphere formation (n=6 biological replicates). Data are displayed relative to control. Representative images are shown in (B). (C) The effects of 0.25μM or 2.5μM cadmium chloride, relative to control, on secondary mammosphere formation (n=6 biological replicates), with representative images are shown in (D).

### Effects of Cadmium Exposure on 3D Branching Organoid Formation

After 7-10 days, primary mammospheres are plated in 3D collagen based hydrogels which mimic the composition of the extracellular matrix of the human breast. Over the course of 2 to 3 weeks, the cells grow into complex branching organoids (Figure 3A), which closely recapitulate mammary gland structures *in situ* in the human mammary gland (Sokol et al. 2016). Unlike in the mammosphere formation experiments, 0.25μM cadmium chloride treatment reduced the number of branching organoid structures formed by approximately 68%, while the 2.5μM exposure led to a reduction of approximately 83%. (Figure 3B). Imaging of hydrogels showed changes to the extent of the branching ductal growth after cadmium exposure (Figure 3C). Cells exposed to 0.25μM cadmium chloride and grown in hydrogels demonstrated ductal growth, but was significantly less pronounced and it did not extend through the hydrogel as extensively as in the control conditions. The 2.5μM cadmium exposed hydrogels had very little growth overall. These data demonstrate that cadmium exposure significantly decreases branching structure formation.

**Figure 3.**
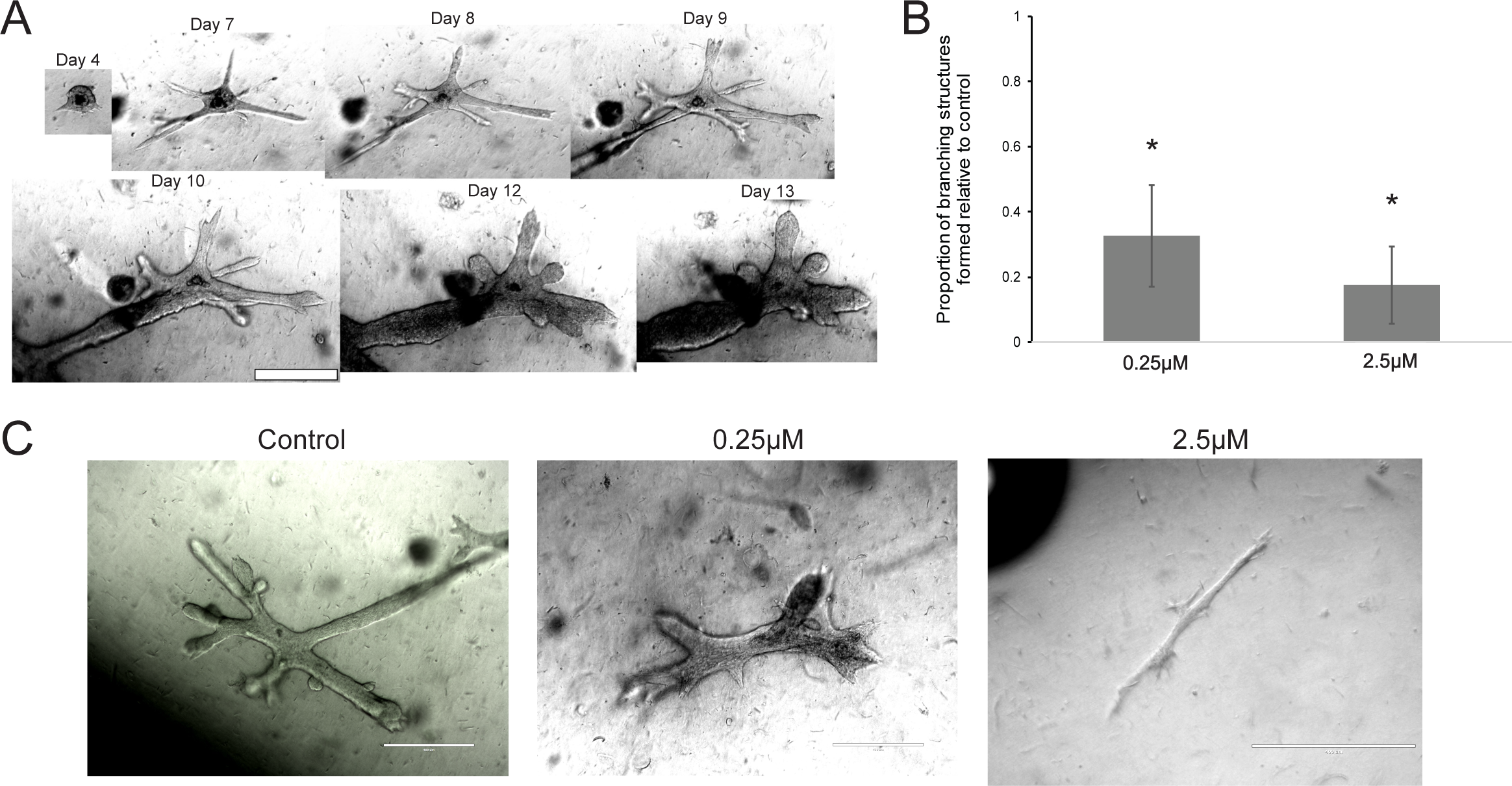
The effects of cadmium on organoid formation. (A) Mammospheres plated in floating collagen-based hydrogels will form complex branching structures over the course of two weeks. (B) The effects of 0.25μM or 2.5μM cadmium chloride on branching structure formation in 3D hydrogels, relative to control (n=3 biological replicates). (C) Representative images of structures formed in each experimental condition.

### RNA-Sequencing of Exposure Treated Primary Mammospheres

The significant decreases in branching morphogenesis caused by cadmium exposure suggested that cadmium treatment was downregulating key developmental pathways involved in stem cell differentiation or invasion. To comprehensively characterize the transcriptional alterations caused by cadmium, RNA was sequenced from 0.25μM and 2.5μM cadmium chloride treated cells, from 4 independent biological replicates, grown in primary mammosphere conditions for 7 days. Primary mammosphere formation yields a population of cells enriched for stem and early progenitor cells (Dontu et al. 2003). Visualization through multidimensional scaling showed that the samples cluster clearly based on individual rather than treatment (Figure 4A). This suggests that the majority of the variance is explained by interindividual differences in breast gene expression. Consistent with the effects on primary and secondary mammosphere formation, 2.5μM cadmium chloride treatment lead to more significant alterations in gene expression compared to 0.25μM treatment (101 and 5 genes altered at FDR < 0.05, respectively, Figure 4B and C; alterations for all genes presented in **Supplemental Tables 1 and 2**). All of the significant changes observed at 0.25μM treatment were also found at 2.5μM treatment (Figure 4D). Hierarchical clustering of the samples based on the expression of genes identified as differentially expressed (FDR<0.10) in the 2.5μM cadmium treatment showed that the 2.5μM cadmium treatment samples clustered distinctly from the control and 0.25μM samples (Figure 4E).

**Figure 4.**
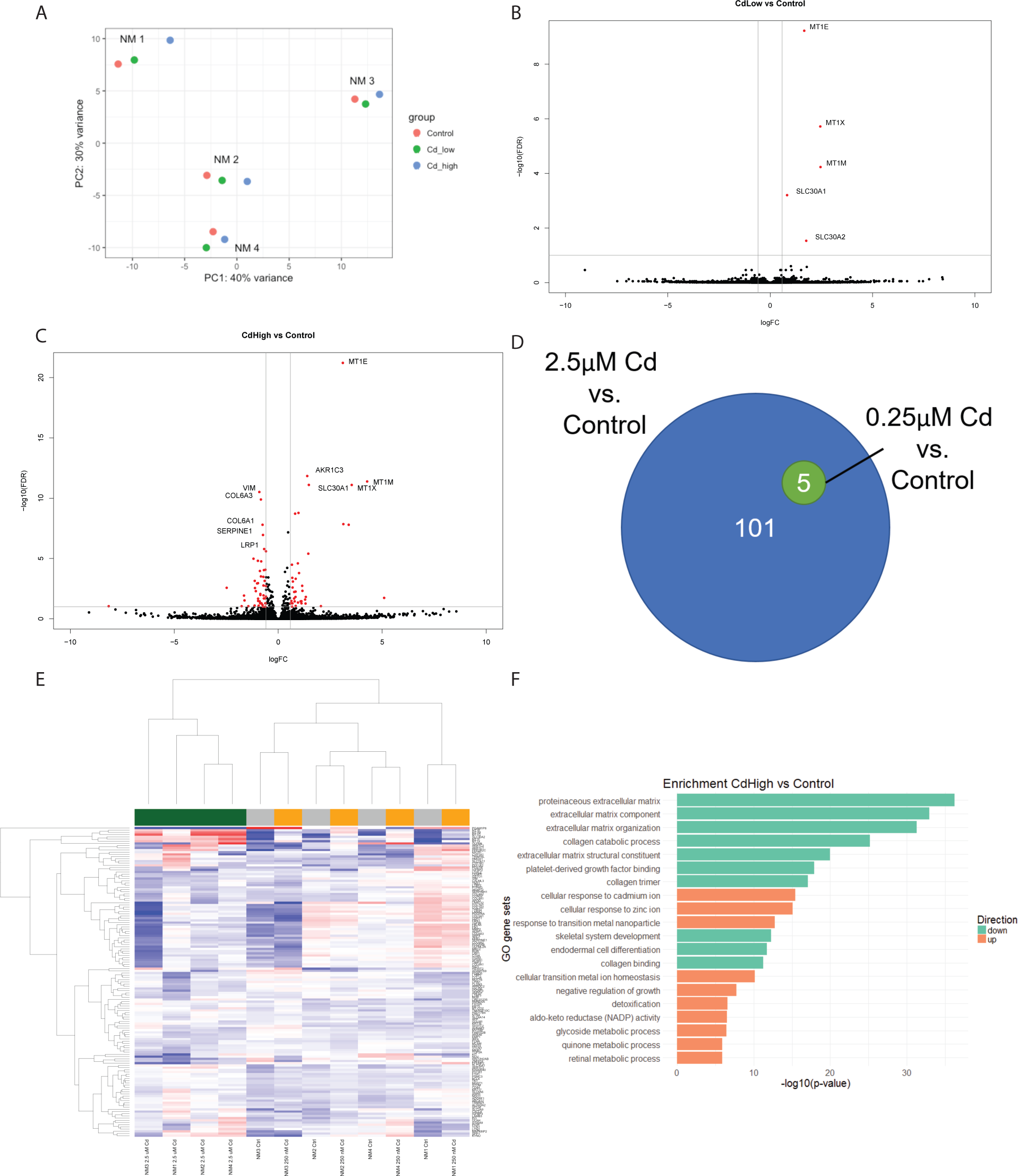
Transcriptomic profiling of the effects of cadmium exposure in primary mammospheres. (A) Multidimensional scaling plot of RNA expression in control, 0.25μM, or 2.5μM cadmium chloride treated mammospheres (n=4 biological replicates). (B) Volcano plot representing differential gene expression in the 0.25μM cadmium treatment vs. control. Red color indicates a significantly differentially expressed gene. (C) Volcano plot representing differential gene expression in the 2.5μM cadmium treatment vs. control. (D) Venn diagram representing the overlap in differentially expressed genes between the 0.25μM cadmium treatment vs. control and 2.5μM cadmium treatment vs. control comparisons. (E) Unbiased clustering analysis of the 12 samples based on expression of the genes identified as differentially expressed between 2.5μM cadmium treatment vs. control. Colorbar: Green = 2.5μM cadmium, Orange = 0.25μM cadmium, Grey = Control. (F) Gene sets enriched for differentially expressed genes in the 2.5μM cadmium treatment vs. control comparison.

### Cadmium Exposure Induces Changes in Metal Response and Extracellular Matrix Related Pathways

At the 0.25μM dose, the 5 genes that were significantly altered were all upregulated, and represent known metal response genes: *MT1E, MT1X,* and *MT1M* are three subtypes of the metal binding protein metallothionein 1 while *SLC30A1* and *SLC30A2* are zinc transporters. These results show that even at low doses of cadmium, stem cell enriched populations of breast epithelial cells are sensitive to the presence of divalent cations and can upregulate biological processes to detoxify or eliminate cadmium. Similar changes were observed at the 2.5μM dose of cadmium chloride, where the top upregulated genes were associated with response to metal ions (Figure 4F). An increase in expression of known oxidative response genes, including *HMOX1, TXNRD1*, and *SPP1*, was also observed. Intriguingly, the pathways most enriched for downregulated genes at the 2.5μM dose were involved in the formation of the extracellular matrix (ECM) or interaction with the ECM (e.g. focal adhesion) (Figure 4F). Specifically, 2.5μM cadmium chloride lead to a decrease in expression of many ECM genes: *COL1A2, COL6A1, COL6A2, COL6A3, FBN1, FN1,* and *NOV.* Further, genes associated with EMT were also downregulated with cadmium treatment, including *ZEB1, VIM,* and *TGFBI.* No changes in known estrogen receptor alpha target genes were noted, including *PGR, C3, GREB1, NRIP1,* or *ABCA3* (Johnson et al. 2003; Lin et al. 2004) (**Supplemental Tables 1 and 2**), suggesting that at these doses and this time point, cadmium is not activating estrogen signaling as a metalloestrogen. Overall, these results show that 2.5μM cadmium chloride exposure increase the expression of metal ion response genes and downregulate the expression of genes involved in ECM production and interaction as well as EMT.

### Cadmium, HIF-1α, and Mammary Stem Cell Growth

A closer examination of the genes and pathways dysregulated by 2.5μM cadmium chloride exposure revealed a consistent downregulation of genes known as targets of the transcription factor hypoxia-inducible factor 1 alpha (HIF-1α). Specifically, we observed a significant downregulation of known HIF-1α targets *LOXL2*, *VIM*, *ZEB1*, *IGFBP2,* and *PDGFRA* (Figure 5A). HIF-1α is required for mammary stem cell expansion and branching morphogenesis (Seagroves et al. 2003). We first tested whether cadmium can inhibit HIF-1α transcription factor activity using a luciferase reporter assay. Both 0.25μM and 2.5μM cadmium chloride significantly reduced the amount of HIF-1α activity (Figure 5B). Additionally, both concentrations of cadmium also significantly attenuated the effects of the pharmaceutical hypoxia mimic and HIF-1α stabilizer FG-4592 ((Beuck et al. 2012); Figure 5B). To test the role that HIF-1α plays in mammary stem cell growth and differentiation, we treated primary mammary epithelial cells with either 0.25μM or 2.5μM cadmium chloride, or 1μM or 5 μM of the pharmaceutical HIF-1α inhibitor acriflavine (Lee et al. 2009). Acriflavine treatment significantly inhibited primary and secondary mammosphere formation and eliminated the formation of any branching organoid structures, a phenotype similar to that observed with cadmium treatment. These results show that HIF-1α activity is important for human mammary stem cell proliferation and branching morphogenesis, and that cadmium, at the doses tested, significantly inhibits the transcriptional activity of HIF-1α.

**Figure 5.**
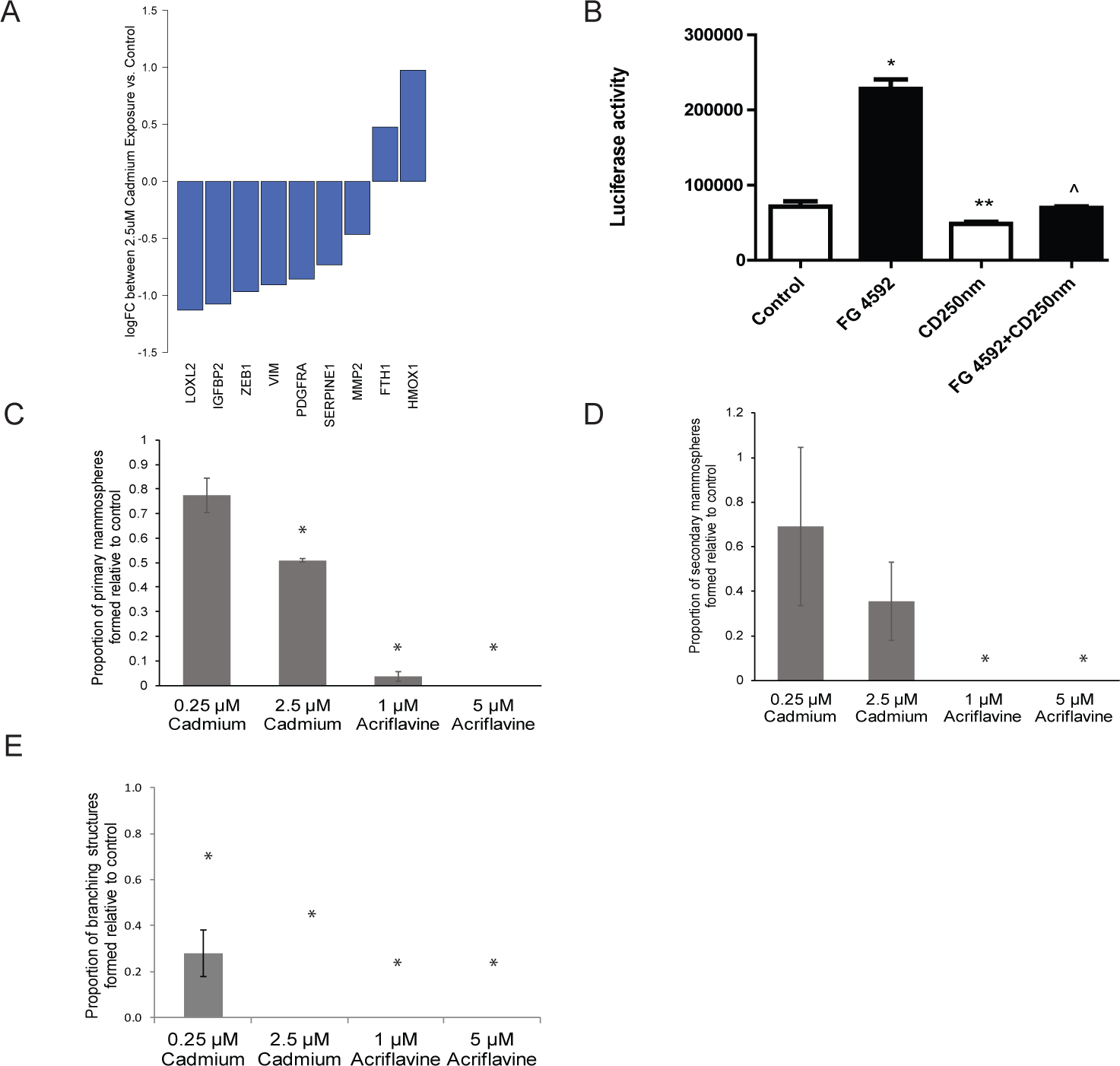
The effects of cadmium on HIF-1α activity and HIF-1α activity on breast stem cell growth. (A) Relative expression of known HIF-1α target genes between 2.5μM cadmium treatment vs. control. (B) Effects of FG 4592, a known HIF-1α activator, 0.25μM cadmium, or a combination of the two on HIF-1α activity as assessed by a luciferase reporter assay. * - p<0.0005 difference from control, ** −p<0.05 difference from control, ^ - p<0.0005 difference between FG 4592 treatment alone. (C) Primary mammosphere formation rates in 0.25μM cadmium, 2.5 μM cadmium, 1 μM acriflavine, or 5 μM acriflavine treatment, relative to control (n=3 biological replicates). (D) Secondary mammosphere formation rates in cadmium or acriflavine treatments relative to control. (n=3 biological replicates) (E) Branching organoid formation rates in cadmium or acriflavine treatments relative to control. (n=3 biological replicates) * - p<0.05 difference from control.

## DISCUSSION

Cadmium is a known human carcinogen, however, its role in human breast carcinogenesis and altered mammary gland development remains controversial. Here, we used 3D tissue culture methods of primary human breast epithelial cells to model the effects of cadmium exposure during times of stem cell expansion and differentiation. We identified that cadmium exposure, at concentrations in the range of those previously detected in human breast tissue, significantly inhibits primary and secondary mammosphere formation, which are functional readouts of stem cell activity and self-renewal capacity (Dontu et al. 2003). Further, cadmium exposure significantly inhibits breast epithelial cell ductal elongation and branching morphogenesis in collagen-based hydrogel cultures which model the human breast ECM (Sokol et al. 2016). To identify pathways and genes dysregulated by cadmium that explain this inhibition of growth, we sequenced RNA from cadmium-exposed primary mammospheres, which are enriched for stem and progenitor cells. Cadmium significantly upregulates genes associated with metal response, while downregulating ECM-production and genes involved in focal adhesion, including known targets of the transcription factor HIF-1α. We validated that cadmium exposure at the doses tested can inhibit HIF-1α activity and showed that pharmacologic HIF-1α inhibition ablates mammary growth in our model. Overall, we present a novel approach to assay the effects of environmental toxicants during breast cancer windows of susceptibility in patient-derived breast cells. We show that cadmium can inhibit mammary gland branching morphogenesis at doses relevant to human exposure and identify HIF-1α dysregulation as an important mechanism of toxicity during mammary gland development.

Studies assaying the effects of cadmium exposure on altered breast development or cancer *in vivo* have identified that the life stage timing of exposure is essential in defining effects. These findings are in line with the breast cancer windows of susceptibility hypothesis, where exposures during key developmental stages, specifically *in utero*, during puberty, and during pregnancy, disproportionately influence mammary gland development and breast cancer risk (Russo 2016). The physiological changes during these developmental windows are driven by stem cells undergoing rapid proliferation and differentiation, which potentially increases their vulnerability to environmental influence. *In vivo* studies of *in utero* exposure show that cadmium alters mammary gland development in a manner consistent with estrogen exposure, leading to an increase in the number of terminal end buds (Davis et al. 2013) (Johnson et al. 2003), a stem cell enriched population which drives adult mammary gland differentiation (Paine and Lewis 2017). Interestingly, however, the increase in terminal end buds was found to be dose dependent, occurring at the lower (0.5μg/kg), rather than higher (5μg/kg) dose tested (Johnson et al. 2003), and were not associated with an increase in DMBA-induced carcinogenesis, unlike estrogen treatment (Davis et al. 2013). Mice exposed to cadmium in the pre-pubertal window, instead, experienced a decrease in the number of mammary gland terminal end buds and decreased overall mammary gland ductal development (Alonso-Gonzalez et al. 2007). Mice treated with cadmium during pregnancy displayed a remodelled mammary gland characterized by an increase in fat content and less active alveolar epithelial cells (Ohrvik et al. 2006). Similarly, exposure of lactating Holstein cows to cadmium lead to a decrease in milk production (Miller et al. 1967). Here, we found that exposure of breast cells derived from adults to low dose cadmium lead to impaired branching morphogenesis. Unlike the studies of cadmium exposure *in utero*, we did not observe an estrogenic effect, as reflected by increased expression of estrogen responsive genes. In aggregate, these results suggest that the timing of cadmium exposure during mammary gland development is likely important, where *in utero* cadmium exposure could lead to an increase in terminal end bud formation, while exposure during puberty or pregnancy could inhibit ductal morphogenesis and proper mammary gland function.

Mechanistic studies of the effects of cadmium on immortalized breast cell lines, both non-tumorigenic and cancer, have identified that cadmium can also induce cancer-related alterations in key developmental processes, including the epithelial-mesenchymal transition (EMT). During EMT, epithelial cells lose their tight junctions to their neighboring cells and gain invasive and migratory properties, which are important for the development of metastatic cancer (Wang and Zhou 2011). A forty-week treatment of the immortalized, but non-tumorigenic, breast cell line MCF10A with 0.25μM cadmium lead to the acquisition of an invasive, aggressive phenotype consistent with basal breast cancers through a mechanism independent of ER-α (Benbrahim-Tallaa et al. 2009). Recently, treatment of MDAMB231 cells, an already aggressive and invasive triple negative breast cancer cell line, with 1uM or 3uM cadmium lead to a more invasive and metastatic phenotype (Wei and Shaikh 2017). While the results of these studies lead us to initially hypothesize that we would observe a similar EMT-like phenotype in cadmium treated primary breast stem cells, we instead observed the opposite. Both our functional and molecular data showed that cadmium treatment of primary cells lead to decreased invasion and growth in hydrogel cultures, further supported by a downregulation of key genes involved in EMT including *VIM* and *ZEB1*. Future research should focus on understanding this unexpected difference in the response of immortalized cell lines and primary breast cells to cadmium, particularly in light of the inconsistent association between cadmium and breast cancer risk in epidemiological studies.

Through our unbiased approach, we found that 0.25μM cadmium exposure lead to the downregulation of multiple known targets of the hypoxia inducible factor HIF-1α. HIF-1α is an oxygen sensing transcription factor that plays an essential role in mammary gland development. For example, while mammary glands from nulliparous targeted HIF-1α knockout mice do not display phenotypic differences from control mice, during pregnancy HIF-1α knockout mice have stunted mammary gland differentiation, ultimately leading to lactation failure (Seagroves et al. 2003). This finding suggests that HIF-1α activity is essential for the mammary gland expansion during pregnancy and lactation, but not mammary gland development *in utero*. This aligns with the findings of the effects of cadmium exposure during puberty and pregnancy significantly disrupting mammary gland formation and leading to different phenotypic effects compared to *in utero* exposure. Using a pharmacologic inhibitor of HIF-1α, acriflavine (Lee et al. 2009), we found that inhibition of HIF-1α ablates mammosphere formation and branching morphogenesis, confirming the importance of this transcription factor in adult mammary gland differentiation. Others have previously reported that cadmium exposure can inhibit HIF-1α transcriptional activity in other organ systems and cell types. Cadmium exposure caused known HIF-1α target genes to not be induced during hypoxia, disrupted HIF-1α DNA binding in Hep3B cells, and decreased luciferase expression in a HIF-1α activity reporter system similar to the one used here (Chun et al. 2000). Others have reported that cadmium exposure in rat lung fibroblasts lead to a decrease in HIF-1α binding to promoter region of the lysyl oxidase gene, a known HIF-1α target (Gao et al. 2013). Taken in aggregate, these results suggest that exposure to cadmium at doses relevant to human exposures can inhibit normal HIF-1α function during human mammary gland differentiation, leading to an inhibition of branching morphogenesis and mammary gland development, particularly during windows of susceptibility such as pregnancy.

The present study has a number of strengths relative to the existing literature. First, this is, to the best of our knowledge, the first study to test the effects of cadmium on patient-derived primary breast cells, rather than immortalized cell lines or in animal models. We developed and used a novel model of human mammary gland development, including a physiologically relevant 3D hydrogel based organoid system (Sokol et al. 2016), to characterize the effects of cadmium exposure during windows of susceptibility. We incorporated an unbiased whole transcriptome analysis to comprehensively quantify the effects of cadmium in a human stem cell enriched population, and functionally validated the findings of HIF-1α inhibition. This study also has a number of weaknesses. We used primary breast cells from adult reduction mammoplasty patients, which may not represent a truly “normal” breast cell population (Degnim et al. 2012), or may not recapitulate the biological effects of cadmium exposure *in utero*. While our cadmium treatments ranged from 7 to 21 days total, we did not attempt to recapitulate the extended cadmium exposure (up to 40 weeks) of other studies, which may better model the effects of chronic exposure to cadmium (Benbrahim-Tallaa et al. 2009). Additionally, for this study, we were blinded to the study participant characteristics besides age, including parity or family history of breast cancer. Thus, we were unable to make any conclusions about epidemiological factors that may predict increased or decreased susceptibility to the effects of cadmium. Future studies incorporating the life stage or other breast cancer risk factors of the donors, while in parallel conducting a longer-term cadmium exposure, may be able to better model human population-relevant effects of cadmium on mammary gland toxicity or cancer risk.

Here, we show that cadmium exposure, at doses relevant to human health, can significantly inhibit human mammary stem cell proliferation and differentiation. These findings are in line with animal studies of cadmium exposure during puberty and pregnancy, whose results stand distinct from studies of *in utero* exposure. The effects observed are likely mediated, at least in part, by cadmium’s disruption of HIF-1α activity, identifying HIF-1α as an important target for human mammary gland developmental toxicity. While our results do not support the hypothesis that cadmium acts as a breast cancer initiator through the induction of stem cell proliferation, our results suggest that cadmium exposure in adult women may significantly inhibit mammary gland proliferation and development.

## Acknowledgements

This work was supported by the Ravitz Family Foundation, a grant P30ES017885 from the National Institute of Environmental Health Sciences (to J.A.C) and CA130810 from National Cancer Institute (to Y.M.S), National Institutes of Health.

## Supplemental Table Labels

**Supplemental Table 1.** Differential expression results from RNA-seq comparing control and 0.25μM cadmium treated mammospheres.

**Supplemental Table 2.** Differential expression results from RNA-seq comparing control and 2.5μM cadmium treated mammospheres.

